# Efficient and specific Ly6G^+^ cell depletion: A change in the current practices toward more relevant functional analyses of neutrophils

**DOI:** 10.1101/498881

**Authors:** Julien Faget, Gael Boivin, Pierre-Benoit Ancey, Aspasia Gkasti, Julie Mussard, Camilla Engblom, Christina Pfirschke, Jessica Vazquez, Nathalie Bendriss-Vermare, Christophe Caux, Marie-Catherine Vozenin, Mikael J. Pittet, Matthias Gunzer, Etienne Meylan

## Abstract

Neutrophils orchestrate the innate immune response against microorganisms and are increasingly recognized to modulate cancer development in primary tumors and metastases. To address their function *in vivo*, different approaches are used, the most common ones relying on antibody-mediated neutrophil depletion. By comparing the effects of two widely used antibodies, we demonstrate a strong efficacy but a lack of specificity for anti-Gr1. In contrast, anti-Ly6G lacks neutrophil-depletion capacity in C57BL/6 mice, which can be explained by an insufficient celerity of neutrophil clearance that is counterbalanced by exacerbated mobilization of immature cells. When combined with a secondary antibody, anti-Ly6G treatment results in specific and efficient neutrophil depletion. Using a mouse model of lung adenocarcinoma, we demonstrate the efficacy of this new approach to diminish primary tumor growth and propose the existence of a local intercellular communication between neutrophils and alveolar macrophages that fosters regulatory T cell proliferation in lung cancer.

## INTRODUCTION

Neutrophils are the most abundant cell type among human leukocytes and play an essential role in host defense against bacteria and in sterile inflammation^1^. They were among the first immune cell types associated with cancer progression^2^ and their involvement in autoimmune diseases such as antibody-associated systemic vasculitis, atherosclerosis, systemic lupus erythematous, rheumatoid arthritis, psoriasis^3–5^ and wound repair^6^ positions them as an essential component in multiple immunological processes. Furthermore, it has recently been shown that neutrophils infiltrate multiple tissues where they instruct homeostatic and pathological conditions by contributing to circadian gene regulation^7^. They are constantly produced and released in the bloodstream by the bone marrow (BM)^8^ and are rapidly cleared within about twelve hours in tissues (spleen, BM and liver) through the action of resident phagocytes^9^. BM neutropoiesis involves about forty percent of BM cells and is tightly regulated through the IL23/IL17/G-CSF axis, itself governed by gut macrophages in physiological conditions^7,10,11^.

Whereas neutrophils can be easily purified from blood and BM, their short half-life, absence of proliferation and high sensitivity to environmental conditions leads to important limitations to study them functionally *in vitro*. Hence, *in vivo* depletion experiments and genetic engineering allowing conditional gene knockouts and reporters are the most relevant approaches to investigate neutrophil biology in various contexts. In mice, neutrophils are characterized by the expression of both Ly6G and Ly6C antigens, with Ly6G being restricted to neutrophils^12^. Until 2008, depletion-based experiments suffered from a lack of specific antibodies (ab) allowing neutrophil targeting without impacting other immune populations. Indeed, the anti-Ly6G/C Gr1 ab (clone RB6-8C5) was extensively used to deplete these cells, but due to its ability to bind both Ly6C and Ly6G antigens, this ab also depletes Ly6C^+^ monocytes and a subset of CD8 T cells, potentially reducing the relevance of the observations. The characterization of the anti-Ly6G specific ab (Clone 1A8) has provided new possibilities, as it only binds neutrophils and seems to be the most relevant agent to target specifically these cells *in vivo^13^*. Genetic approaches based on LysM-Cre^14,15^ or MRP8/S100A8-Cre^16^ were developed to manipulate and trace neutrophils *in vivo* but the transgenes are also expressed at least transiently by other myeloid lineages when bred with ROSA-EYFP (Gt(ROSA)26Sor^tm1(EYFP)Cos^) animals^17^. Recently, the knockin of a bi-cistronic sequence coding for the Cre recombinase and the fluorescent protein tdTomato into the *Ly6g* locus (designated as Catchup mice) led to specific expression of Cre in neutrophils, allowing both genetic manipulation and cell tracing to an extent that had never been reached before. Indeed, breeding of these mice into a strain carrying the Cre-activatable tdTomato reporter under the CAG promoter into the *ROSA26* locus, giving rise to the Catchup^IVM-red^ strain, demonstrated specific conditional expression of the tdTomato in neutrophils^18,19^. Here, we have used this mouse strain to generate mice having a conditional expression of the diphtheria toxin receptor (DTR) and evaluated them in neutrophil depletion experiments.

We recently demonstrated that neutrophils are the main contributors of disease progression among tumor infiltrating immune cells in the *Kras^G12D/WT^; p53^fl/fl^* inducible model of autochthonous lung cancer (KP mice)^20,21^. In these publications, neutrophil tumor promoting activity was assessed through depletion experiments based on anti-Gr1 ab (clone RB6-8C5) as the anti-Ly6G (clone 1A8) failed to deplete these cells. In the present study, we investigated the limitations of these abs observed in different research centers, identified a new method to achieve specific and efficient neutrophil depletion, and further used this method based on ab combination to reveal the impact of neutrophils on lung tumor microenvironment and progression.

## Results

### The anti-Ly6G ab clone 1A8 does not deplete neutrophils in C57BL/6 mice

In surveying the scientific literature over a one-year period we found forty-five publications using the key words “(Neutrophils and (depletion))” on PubMed, in which the authors reported efficient neutrophil depletion. One-third of these publications were using anti-Gr1 ab to remove neutrophils *in vivo*, whereas two-thirds used anti-Ly6G ab (figure 1a). To assess whether these abs indeed deplete neutrophils, we investigated their effects in 14 weeks old Catchup^IVM-red^ (C57BL/6) mice whose neutrophils can be easily and specifically identified based on tdTomato expression. We found that anti-Gr1 ab treatment strongly reduced the proportion of tdTomato positive neutrophils; however, the percentage of neutrophils remained unchanged upon anti-Ly6G treatment (figure 1b). To address if mouse strain, origin (produced in our facility or not), housing conditions or age could affect anti-Ly6G depletion activity, we compared mice of three different strains, C57BL/6J, BALB/c and FVB/N imported from Charles River, and treated them at different age: 9-weeks old (housed for two weeks in our facility) and 24-weeks old (housed for 17 weeks). In these experiments, we identified circulating neutrophils using flow cytometry as CD11b^+^, CD45^+^, Ly6C^int^ cells. All strains presented a proportion of neutrophils of 7.6% ± 2.1 among immune cells at nine weeks of age and anti-Ly6G treatment (200μg) led to neutrophil depletion reaching 71%, 90% and 95% in C57BL6/J, BALB/c and FVB/N strains respectively, one day after ab injection. In 24-weeks old mice (housed in our animal facility for 17 weeks) all strains showed a higher proportion of circulating neutrophils (14.3% ± 3.8) when compared to the same strain at 9-weeks of age and housed in our facility over two weeks. Depletion with anti-Ly6G treatment and using the same protocol was ineffective in C57BL/6J mice (average neutrophil proportion was 187% of the controls) whereas it remained efficient in BALB/c (92.7%) and FVB/N (84.31%) mice (figure 1c). These observations raised the possibility that age and/or housing conditions contribute to the resistance of C57BL/6J mice to neutrophil depletion. To address this, we imported 18-weeks old C57BL/6J mice from Charles River and treated them with anti-Ly6G ab after two weeks of acclimation in our mouse facility. In these mice, the proportion of circulating neutrophils was similar or lower in controls to what we observed in nine-weeks old mice (4.6%± 1.2). Nevertheless, the injection of 200μg of anti-Ly6G ab failed to change the proportion of neutrophils among immune cells (figure 1d). Hence, these observations suggest that neutrophil proportions in the circulation depend on the housing conditions in all strains of mice and that the anti-Ly6G ab is largely ineffective at depleting neutrophils in C57BL/6J mice.

**Figure 1:**
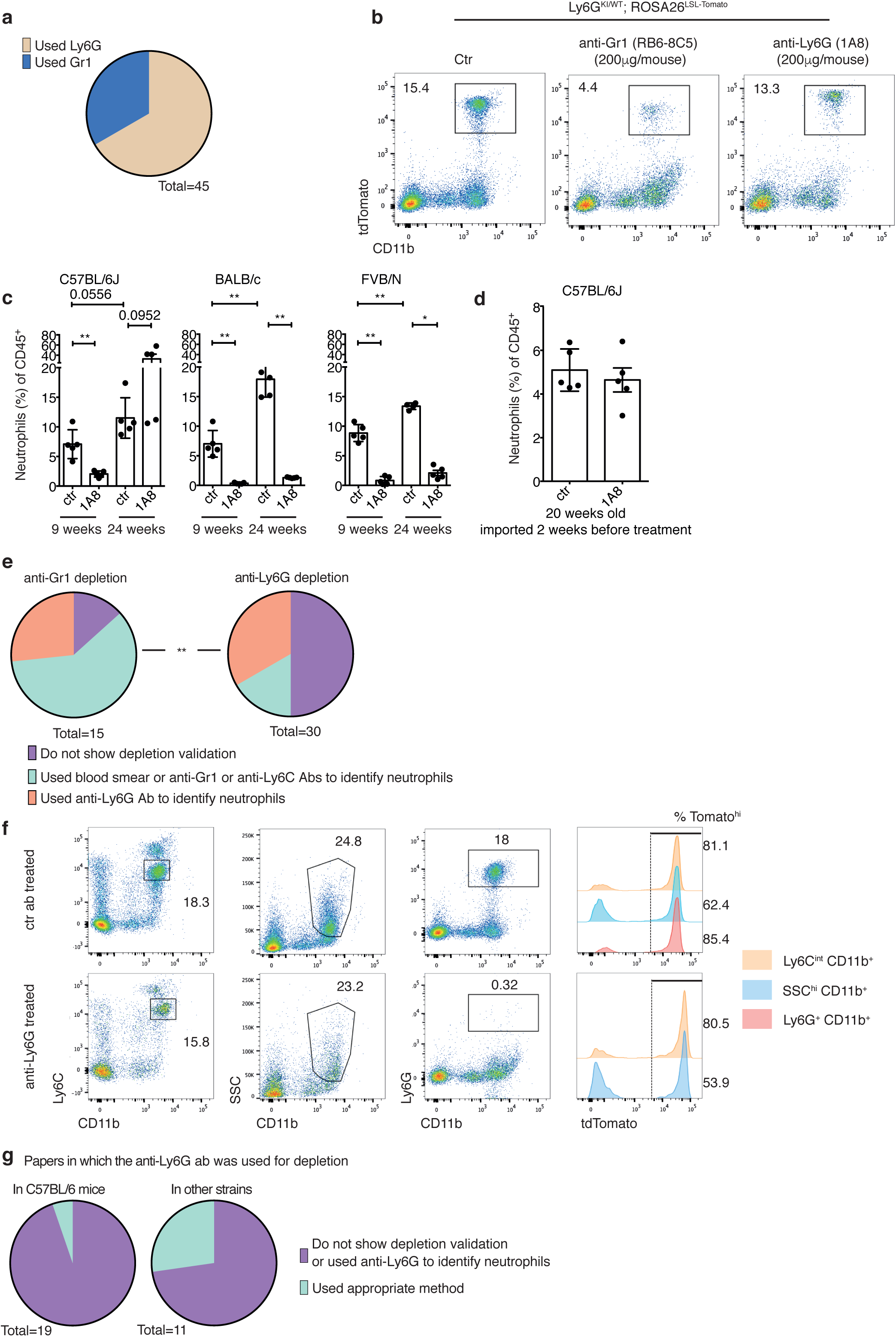
Anti-Gr1 (RB6-8C5) but not anti-Ly6G (1A8) ab allows efficient depletion of neutrophil in C57BL/6J mice. a) Analysis of the literature (April 2017 to March 2018, total = 45 studies). b) Flow cytometry plots showing neutrophil proportion identified as CD11b^+^tdTomato^+^ cells among CD45^+^ cells in 14 weeks old Catchup^IVM^-^red^ mice treated with 200μg of control ab (Ctr: clone 2A3), anti-Gr1 or anti-Ly6G ab, 24 hours before sampling. c) Percentage of neutrophils, identified as CD45^+^CD11b^+^Ly6C^int^. C57BL/6J, BALB/c and FVB/N mice imported from Charles River and treated with anti-Ly6G or control (ctr) ab (200μg) for 24 hours at 9 weeks and 24 weeks of age, and housed respectively during 2 or 17 weeks in the EPFL animal facility. n=4-5 mice per condition. d) 18 weeks old C57BL/6J mice from Charles River then housed for two weeks in the EPFL animal facility, then treated as in c. b, c and d) ** p<0.01 from Mann-Whitney test, error bars represent SEM. e) Strategies used in papers identified in (a) to demonstrate neutrophil depletion efficacy when anti-Gr1 (left) or anti-Ly6G (right) abs were used. ** p<0.01 using Chi-scare test. f) Flow cytometry plots showing gating strategies to identify neutrophils in Catchup^IVM-red^ mice treated as in a). Percentages of tdTomato^+^ cells among gated populations are provided with the histograms. g) Proportion of publications using anti-Ly6G ab to deplete neutrophils in C57BL/6J mice (left, n=19) and other strains (right, n=11).

Confirming our observations, results from two other Institutions (MGH, USA; CRCL/CLB, France) showed that depletion using anti-Ly6G ab was not effective in C57BL/6J mice following different treatment modalities (figure S1a and b).

Considering the important number of publications reporting anti-Ly6G ab based depletion of neutrophils in C57BL/6 mice, we decided to investigate what could be the main differences between our practice and what we can find in the literature. In the majority of publications using anti-Gr1, the validation of neutrophil depletion was done based on CD11b/Ly6C staining or blood smears. On the other hand, from thirty articles published the same year and using anti-Ly6G ab, ten identified neutrophils based on Ly6G and CD11b expression and only five provided evidence of neutrophil depletion using blood smears or CD11b/Ly6C staining (figure 1e). This prompted us to test different gating strategies based on anti-Ly6G, anti-CD11b and anti-Ly6C co-staining on blood cells from Catchup^IVM-red^ mice treated one day before with 200μg of anti-Ly6G ab (clone 1A8) or control ab. We observed that Ly6C^int^/CD11b^+^ cells correspond well to the neutrophil population as 80% and 81% of these cells were expressing the tdTomato in both control and anti-Ly6G treated conditions, respectively. Alternatively, gating on neutrophils based on their granularity (side scatter high) and CD11b also contained non-neutrophil cells as only 62% and 53% of the gated cells were tdTomato positive, respectively in control and Ly6G ab treated mice. Importantly, we failed to detect any cells in the Ly6G^hi^/CD11b^+^ gate in mice treated with the anti-Ly6G ab, demonstrating that masking or internalization of the Ly6G antigen occurs when mice are treated with anti-Ly6G (figure 1f). We performed a titration of the anti-Gr1, anti-Ly6G and anti-Ly6C antibodies on blood samples from Catchup^IVM-red^ mice treated with either anti-Ly6G, anti-Gr1 or control abs (200μg/mouse 24 hours before blood sampling). Strikingly, in both anti-Gr1 and anti-Ly6G treated animals, staining of neutrophils (identified as tdTomato positive cells) is impossible with the anti-Ly6G ab. The anti-Gr1 ab gives a weak signal, which could be used to identify neutrophils when mice are treated with anti-Ly6G, and anti-Ly6C staining is not impacted by any of the treatments (figure S1c). Hence, the strategy that is used to monitor neutrophil depletion in the majority of papers using the anti-Ly6G ab is likely to be misleading. Finally, supporting the fact that anti-Ly6G treatment is most probably not effective in C57BL/6J mice, we observed that, over twenty articles using anti-Ly6G in that strain, only one provides valid data to evaluate neutrophil depletion but in this case, authors failed, in these correct settings, to show any significant impact on neutrophils in 10 to 13-weeks old mice (figure 1g).

### Celerity of neutrophil clearance is the limiting factor to achieve neutrophil depletion

We demonstrated that treatment of C57BL/6J mice with anti-Ly6G ab ineffectively reduces the number of circulating neutrophils but, according to the literature, it is also clear that treating mice with this ab has an effect on neutrophil biology. Hence, we hypothesized that anti-Ly6G treatment leads to a qualitative change of circulating neutrophils. We treated C57BL/6J mice with anti-Ly6G or anti-Gr1 ab, then compared the remaining neutrophils to the control condition using Giemsa staining on blood smears. This demonstrated that both anti-Ly6G and anti-Gr1 abs lead to the replacement, in the circulation, of mature neutrophils showing full nuclear segmentation by immature cells displaying band C shape or bisegmented nuclei (figure 2a). Hence, both anti-Ly6G and anti-Gr1 abs are able to increase neutrophil renewal but the anti-Ly6G ab is less efficient as it only leads to a qualitative modification of neutrophils and fails to achieve a quantitative decrease of the circulating cells. These findings suggest that anti-Ly6G treatment inefficiently depletes neutrophils due to rapid mobilization of immature neutrophils from the BM to the periphery. The anti-Ly6G ab is a rat-IgG2a isotype orthologous to the mouse IgG1 heavy chain, whereas the anti-Gr1 ab is a rat-IgG2b isotype orthologous to the mouse IgG2a heavy chain^22^. We reasoned that this difference could explain why only the anti-Gr1 ab can decrease the number of circulating neutrophils. To investigate this, we combined the anti-Ly6G ab with a mouse IgG2a ab recognizing the rat IgGκ light chain (clone MAR 18.5 that will bind to the anti-Ly6G ab). To do so, we sequentially treated 12 weeks old Catchup^IVM-red^ mice with anti-Ly6G (n=9) or control ab (n=4) (day one) followed by anti-rat injection (day two) for every mouse except four of the ones that received anti-Ly6G only. Monitoring of tdTomato^+^ cells at day 3 revealed a complete depletion of neutrophils in the combination treatment (Ly6G plus anti-rat) but not in controls or when anti-Ly6G was given alone. Furthermore, the Ly6G epitope was hidden by treatment in all conditions where the mice were injected with anti-Ly6G, confirming the binding of anti-Ly6G ab on circulating neutrophils *in vivo* (figure 2b). We then investigated if anti-Ly6G alone or in combination with anti-rat-IgGκ exacerbates neutropoiesis. We found that Lineage^−^ (Lin^−^), Sca-1^+^cKit^+^ hematopoietic stem cells (LSK cells) were increased in number and proportion upon both treatments (figure 2c). Similarly, the two treatment modalities also led to an increase of the proportion and the number of common monocyte/granulocyte precursors (GMP) identified as Lin^−^cKit^+^CD41^Low^CD150^−^CD16/32^hi^ (figure 2d and S2a). Accordingly, we noticed that mature neutrophils identified as s100A9^+^Ly6C^+^CD62L^+^CXCR2^+^ cells totally disappeared from BMs of animals treated with either anti-Ly6G or the combination (figure S2b).

**Figure 2:**
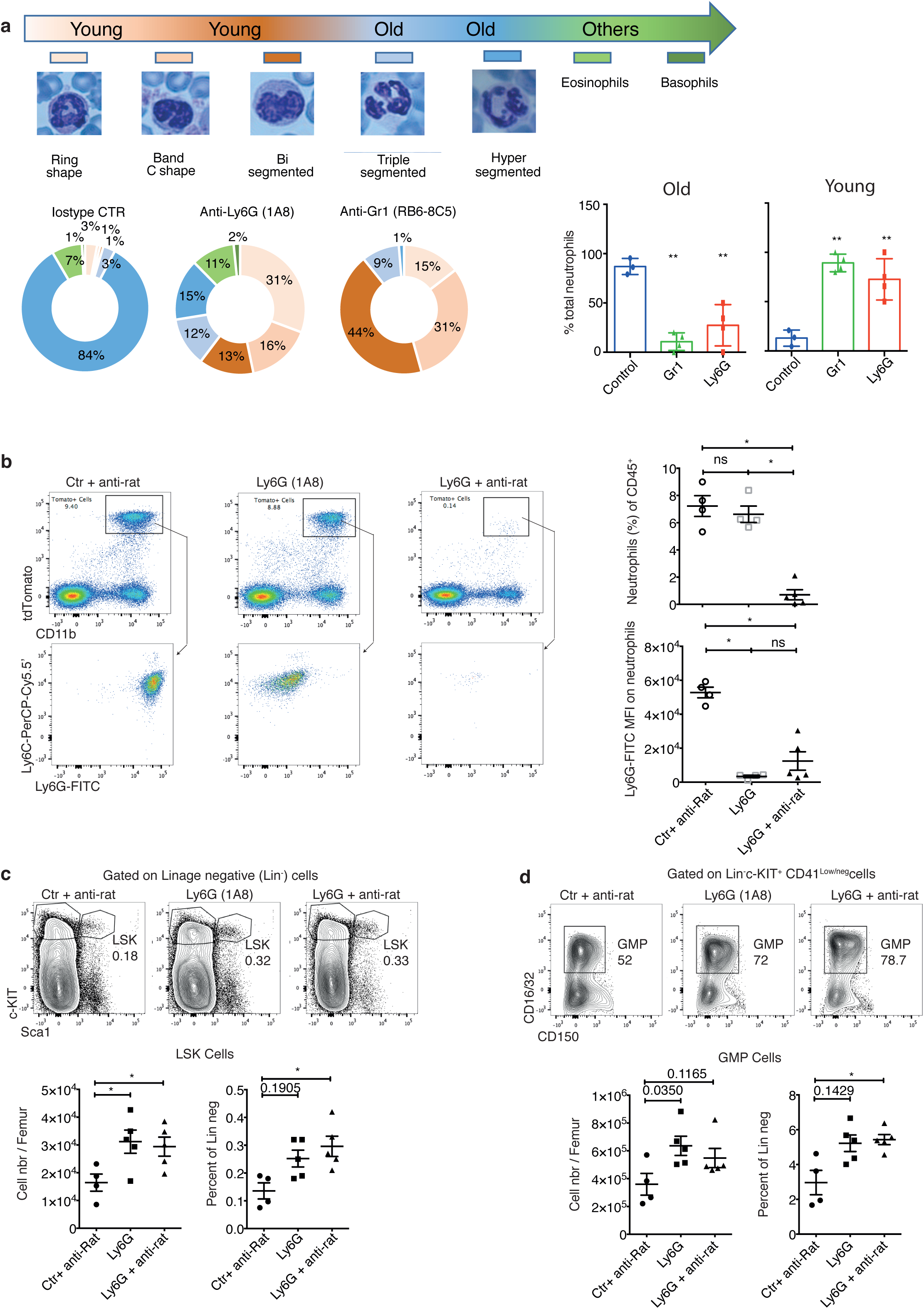
Combination of anti-Ly6G plus anti-Rat-IgGκ abs leads to profound neutrophil depletion in C57BL/6J mice. a) Analysis of neutrophil age based on their nuclear morphology on blood smears 24h after treatment with 100μg of anti-Ly6G or anti-Gr1 ab. Neutrophils were classified as young when they displayed ring-shape, band-C shape and bi-segmented nuclei. Cells showing triple-segmented and hyper-segmented nuclei were assigned as old neutrophils. Histograms show the percentage of young and old cells among total neutrophils (Ctr, n=3, Ly6G and Gr1 n=4). b) Flow cytometry plots showing (top) the percentage of neutrophils gated as CD11b^+^tdTomato^+^ cells in Catchup^IVM^-^red^ and (bottom) results of Ly6C and Ly6G staining on blood samples from mice treated at day one with 100μg of anti-Ly6G or control ab (Ctr) then 100μg of anti-Rat-IgGκ (clone MAR18.5) at day two (^+^ anti-rat). Blood sampling and analysis were done at day three. Histograms show (top) percentage of CD11b^+^tdTomato^+^ neutrophils and (bottom) MFI of Ly6G-FITC ab on these cells. (n=4 for ctr ^+^ anti-rat and anti-Ly6G alone, n=5 for anti-Ly6G ^+^ anti-rat). c-d) Flow cytometry plots showing the gating strategy used to identify c) LSK or d) GMP hematopoietic stem cells and progenitors in bone marrow from C57BL/6J mice treated as in c). Histograms give the number per femur (left) and percentage (right) of the cells of interest based on true-count flow cytometry. n=4-5, a-d) * p<0.05, ** p<0.01 from Mann-Whitney test, error bars represent SD a) and SEM b-d).

Altogether, our results demonstrate that, contrasting to anti-Gr1, the anti-Ly6G ab fails to reduce quantitatively circulating neutrophils in C57BL/6J mice but modifies them qualitatively by increasing neutrophil renewal in peripheral blood. This is probably because the anti-Ly6G ab is a rat-IgG2a isotype orthologous of mouse IgG1. Hence, to achieve a complete depletion of these cells using the anti-Ly6G ab it is mandatory to combine it with an anti-rat-IgGκ mouse IgG2a secondary ab.

### Combining anti-Ly6G plus anti-rat IgGκ abs achieves specific neutrophil depletion

Already existing transgenic mice such as the LysM-Cre^14^ and MRP8-Cre^16^, when bred with ROSA-EYFP (Gt(ROSA)26Sortm1(EYFP)Cos) mice, display approximately 80% of neutrophils among blood circulating YFP positive cells^17^. Hence, with the aim to establish a model where neutrophils can be specifically depleted in an inducible manner and over a long time period without using abs, we decided to cross the Catchup mice with the iDTR strain^23^, where a loxP-flanked STOP cassette governs the expression of the simian DTR gene (HB-EGF) knocked into the Gt(ROSA)26Sor locus, designated as Catchup^DTR^ mice. To compare abs and this genetic approach to deplete neutrophils, we treated 12-weeks old mice over nine days with control plus anti-rat-IgGκ abs, anti-Ly6G plus anti-rat-IgGκ abs, anti-Gr1 ab (until day five) and diphtheria toxin (DT, 200μg/kg every two-three days for Catchup^DTR^ mice) (figure 3a). To refine our assessment of the abundance of neutrophils and other populations in peripheral blood, we decided to use true-count flow cytometry to monitor the number of circulating immune cells (CD45^+^) and among them: neutrophils (Ly6C^int^CD11b^+^), monocytes (Ly6C^hi^CD11b^+^), CD8^+^ T cells (CD3^+^CD8^+^), Ly6C^+^ CD8 T cells (CD3^+^CD8^+^Ly6C^+^) and CD4^+^ T cells (CD3^+^CD4^+^). At day 0, numbers of total immune cells (3.2±1.2×10^6^ cells), neutrophils (1.5 ± 0.6 ×10^5^ cells), monocytes (4.3±2.8 ×10^4^ cells) and Ly6C^+^CD8 T cells (5.1 ±1.6 ×10^4^ cells) per ml of blood presented with an important inter-mouse variability but were never significantly different between treatment and control groups. Specifically, Catchup^DTR^ mice showed a high number of circulating immune cells compared to normal C57BL/6J mice used for ab treatment (4.7±1.1×10^6^ versus 2.6±0.7×10^6^ cells, respectively in Catchup^DTR^ and wild type C57BL/6J mice). Whereas the number of neutrophils was equivalent between these strains (1.6±0.8×10^5^ and 1.5±0.6×10^5^ cells), the number of monocytes (7.3±2.8×10^4^ and 3.2±1.7×10^4^ cells) and Ly6C^+^ CD8 T cells (6.8±1.6×10^4^ and 4.5±1×10^4^ cells) were higher in Catchup^DTR^ mice than in wild type C57BL/6J animals. From that we concluded that comparison of our different depleting strategies was relevant only in regard to their specific control and not across mouse strains. We observed that mice treated with anti-Ly6G plus anti-rat-IgGκ abs showed efficient, specific and long-lasting neutrophil depletion (day two: 88.1%, day five: 94.1% and day nine: 85.1%). As expected, anti-Gr1 ab treatment was highly efficient in depleting neutrophils over the treatment duration (day two: 98.4% and day five: 98.2%), followed by a restoration of neutrophil number at day nine (149%), four days after treatment interruption. Consistent with the fact that anti-Gr1 ab also binds to Ly6C, we observed a depletion of monocytes (95.4% and 83.3%) and Ly6C^+^ CD8^+^ T cells (60.8% and 76.3%) at days two and five. Four days after treatment interruption (day nine), like for neutrophils, the number of monocytes went back to normal levels whereas Ly6C^+^CD8^+^ T cells remained depleted at 75.9%, demonstrating the low renewal capacity of this lymphocyte subtype in comparison to neutrophils and monocytes. Surprisingly, neutrophil depletion in Catchup^DTR^ mice treated with DT did not work. To confirm that neutrophils are resistant to DT, we purified these cells, validated HB-EGF/DTR expression (figure S3a) and treated them *in vitro* with low to extremely high doses of DT. Supporting our *in vivo* observations, we did not observe any impact of DT on cell viability (figure S3b). As an internal control validating our experimental settings, the number of circulating CD4^+^ T cells was not impacted significantly by any treatment (figure 3b). Hence, our results demonstrate the efficacy and specificity of the ab combination approach and also suggest that DT-based strategies do not always work to deplete neutrophils, supporting the need to improve ab-based depletion strategies.

**Figure 3:**
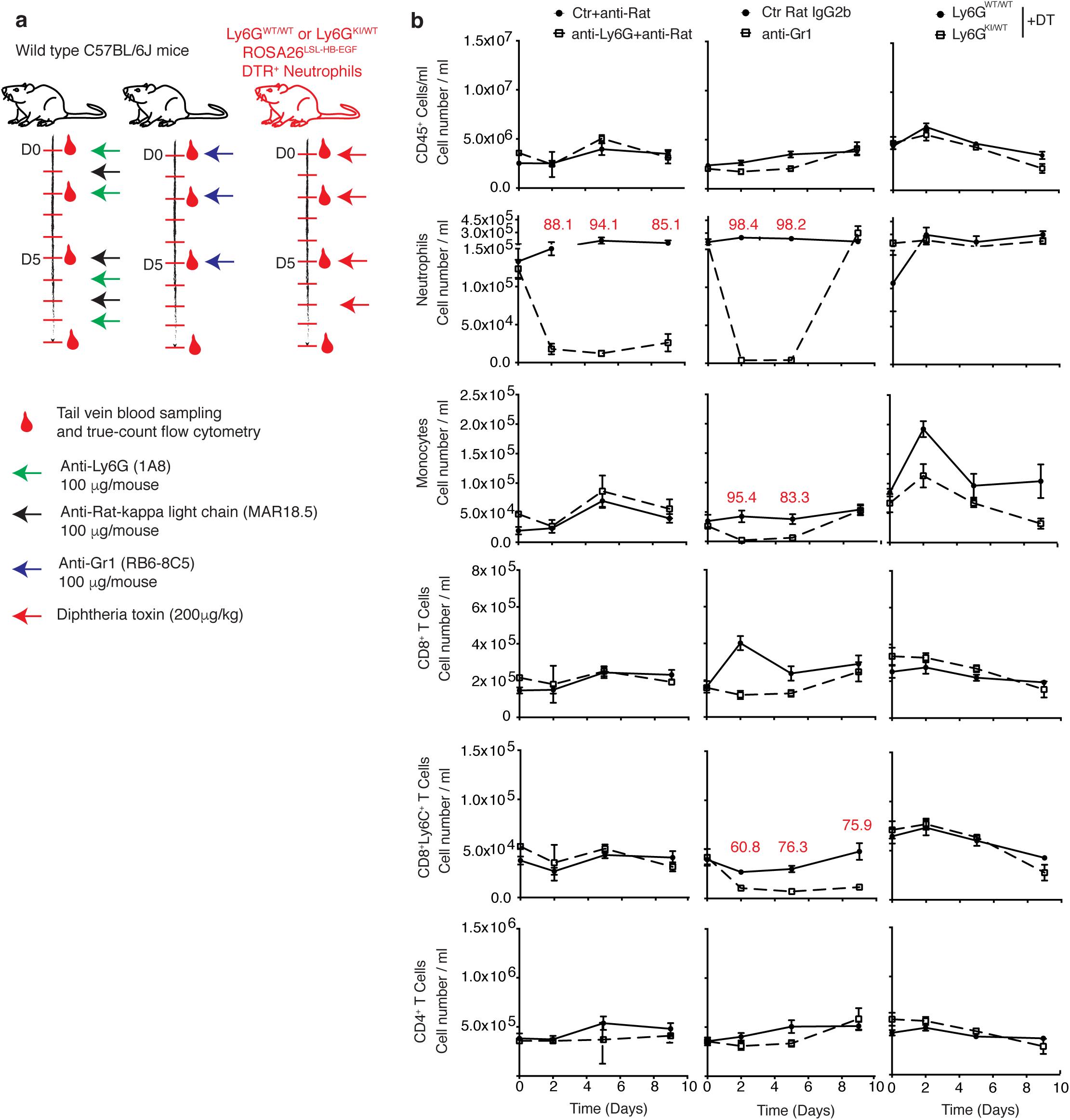
Combination of anti-Ly6G plus anti-Rat-IgGκ abs allows specific and stable neutrophil depletion. a) Schematic view of the experimental plan, six mice per group were treated with control or anti-Ly6G plus anti-Rat-IgGκ abs or control plus anti-Gr1 ab (100 μg per injection). Six Catchup^DTR^ (Ly6G^KI/WT^; ROSA26^LSL-HB-EGF^) mice or littermate controls (Ly6G^WT/WT^; ROSA26^LSL-HB-EGF^) received 200μg/kg of DT every two days. Anti-Gr1 or respective control treatments were stopped at day five. True-count flow analyses were done on 30μl of tail vein blood samples without red blood cell clearing on every mouse at each indicated time point. b) Curves indicate numbers of total immune cells (CD45^+^), neutrophils (Ly6C^int^CD11b^+^), monocytes (Ly6C^hi^CD11b^+^), CD8^+^ T cells (CD3^+^CD8^+^), Ly6C^+^ CD8 T cells (CD3^+^CD8^+^Ly6C^+^) and CD4^+^ T cells (CD3^+^CD4^+^). Red numbers give average depletion efficiency in percentage when compared to respective control. Error bars indicate SEM.

To challenge our approach based on anti-Ly6G plus anti-rat-IgGk, we treated mice over a two-weeks period and observed that in this setting neutrophil depletion loses its efficacy over time (figure S3c). By analyzing Ly6G staining in mice receiving only the anti-Ly6G ab and the mice treated with the ab combination, we observed that the Ly6G epitope was accessible for anti-Ly6G staining only in escaper mice (mice where neutrophil proportion was above 30% of the control at the end of the experiment) from the combination group. In other words, neutrophils in escaper mice are not fully covered by the anti-Ly6G ab injected *in vivo* hence flow cytometry Ly6G staining works on these cells. This suggests that the anti-rat IgGκ antibody can accumulate in the serum of mice and neutralize the anti-Ly6G ab injected to target neutrophils (figure S3d). Accordingly, a positive correlation between the number of neutrophils and the mean fluorescence intensity (MFI) on remaining neutrophils of the anti-Ly6G-FITC ab used for flow cytometry was only observed in mice treated with the combination (figure S3e). Thus, a regular monitoring of Ly6G coverage by anti-Ly6G ab during treatment is critical to adjust the relative concentrations of each ab in order to sustain the efficient depletion for a long time period (see supplementary method part a).

### Alveolar and interstitial macrophages are modulated by tumor-infiltrated neutrophils in lung cancer

To functionally interrogate the relevance of our new approach to deplete neutrophils we tested it on KP mice bearing well-established lung tumors. Similarly to our previous observations where neutrophils were depleted using anti-Gr1 ab^20,21^, mice treated with anti-Ly6G plus anti-rat IgGκ abs demonstrated a significant reduction of tumor growth as monitored using longitudinal micro-computed tomography (μCT) over twelve days of treatment. This result contrasted with the fact that treating KP mice with anti-Ly6G ab alone did not impact tumor growth over the same duration (figure 4a). Hence, increasing neutrophil renewal in the peripheral blood without reducing their number, achieved by anti-Ly6G treatment, fails to abrogate their tumor promoting activity. We then extracted the immune signature from resected tumors using seventeen colors flow cytometry (see supplementary method part b) and we confirmed that anti-Ly6G ab alone did not change the proportions of neutrophils (CD11b^+^S100A9^+^Ly6C^+^) in the tumor mass. Furthermore, despite our efforts and the fact that the combination condition profoundly reduced the proportion of circulating neutrophils at day 11 (figure S4a), we only achieved a 50% reduction of the proportion of neutrophils among tumor immune cells. This demonstrates that even in optimized settings depleting tumor neutrophils remains challenging (figure 4c).

**Figure 4:**
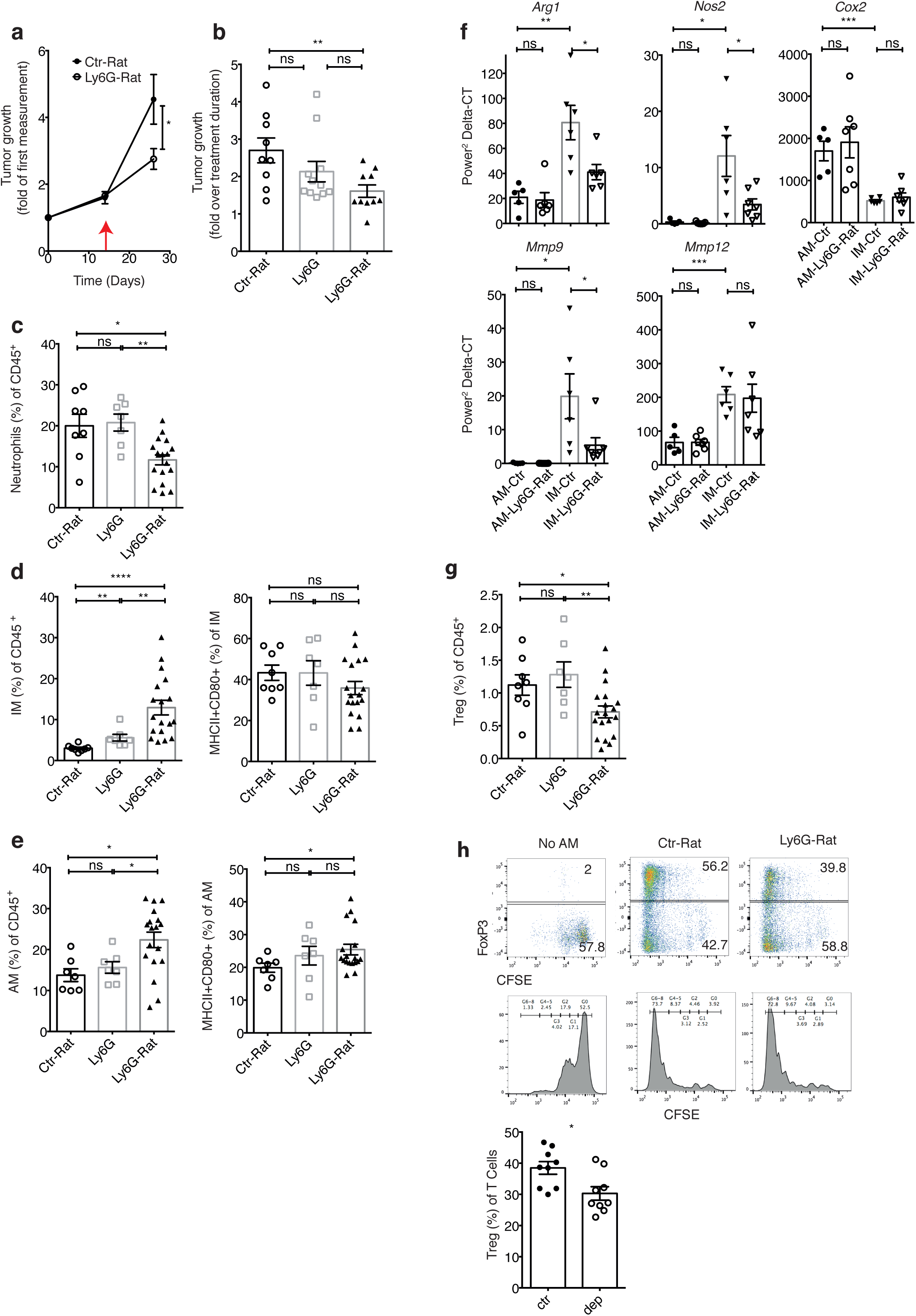
Combination of anti-Ly6G plus anti-Rat-IgGκ abs reduces tumor growth and modifies tumor macrophage recruitment and behavior. a) KP mice were treated with control (ctr) or anti-Ly6G ab (200μg / injection) plus anti-rat lgG_κ_ (100μg / injection) over 12 days. Curves show tumor volume evolution monitored by μCT, red arrow indicates treatment-starting date. b) KP tumor growth rate over 12 days of treatment with control ab plus anti rat-IgGκ (Ctr-Rat), anti-Ly6G alone (Ly6G) or anti-Ly6G plus anti-rat-IgGκ (Ly6G-rat) abs as in a). c-e) Results obtained from 17 colors flow cytometry in KP tumors. c) Percentage of neutrophils (s100A9^+^Ly6C^int^CD11b^+^). d, e) (left) Percentage of d) CD11b^+^F4/80^+^SiglecF^−^ (interstitial macrophages, IM) or e) CD11b^−^F4/80^+^SiglecF^+^ (alveolar macrophages, AM) and double positive CD80^+^MHCII^+^ cells among them (right). f) Relative mRNA expression of the indicated genes on sorted AM and IM from KP tumors. n=5-7. g) Percentage of Treg (FoxP3^+^CD4^+^CD3^+^) from KP tumors. h) Purified syngeneic CD4^+^ T cells stained with CFSE at D0 were cultivated for six days with sub-optimal quantity of T cell expander beads (1 bead for 10 T cells) with or without AM purified form resected tumors of KP mice treated with ctr ^+^ anti-rat or Ly6G ^+^ anti-rat abs as in a). Cells were then stained with anti-CD3 and anti-Foxp3 abs for flow cytometry analysis. Histogram gives the percentage of Foxp3^+^ T cells (n=9). c-e, g) Ctr + anti-rat (n=8), Ly6G (n=7) or anti-Ly6G ^+^ anti-rat (n=17). Error bars indicate SEM, * p<0.05, ** p<0.01, **** p<0.0001 from Mann-Whitney test.

Besides neutrophils themselves, among all the immune populations that we monitored, the most striking effect of neutrophil depletion was an increase of both CD11b^+^SiglecF-F4/80^+^ interstitial macrophages (IM) and CD11b-SiglecF^+^F4/80^+^ alveolar macrophages (AM) in the tumor mass. This latter population showed an increase in the proportion of cells displaying an activated (MHCII^+^ and CD80^+^) phenotype (figure 4d and e). Importantly, these results recapitulated our previous data obtained with the anti-Gr1 Ab (figure S4b). To better characterize potential modifications of tumor macrophage functions upon neutrophil depletion we performed real time PCR analyses of genes known to be associated with macrophage polarization and lung cancer progression on sorted alveolar and interstitial macrophages. Genes linked with lung cancer progression such as *Arg1, Nos2* and *Mmp9* were down-regulated in IM upon neutrophil depletion. On the other hand, *Mmp12, Siglec5* (SiglecF), *Cd274* and *Il10* expression was not different (figure 4f and S4c).

The immune signature also revealed a marked reduction of the regulatory CD4^+^ T cell (Treg: CD3^+^CD4^+^Foxp3^+^) proportion and their proliferation based on Ki67 staining (figure 4g and S4d). This observation was consistent with results obtained using anti-Gr1 ab showing a decrease in Treg proportion among CD4^+^ T cells in tumors (figure S4e). AMs are produced from fetal liver hematopoiesis^24^ and are known to support Treg polarization in physiological conditions^25,26^. Hence, we decided to cell-sort AMs from control and neutrophil depleted tumors to compare their ability to induce Treg enrichment *in vitro*. AMs from tumors supported total CD4^+^ T cell amplification and activated the proliferation of FoxP3^+^ cells. Interestingly, AMs sorted from neutrophil-depleted tumors were also able to support T cell amplification but the resulting T cell pool showed a reduced proportion of FoxP3^+^ Treg (figure 4h).

Altogether, these results suggest that neutrophils interfere with AM function in lung tumors, ultimately supporting their ability induce Treg enrichment.

## Discussion

This technical report evaluates current practices allowing neutrophil depletion and proposes an optimized protocol to achieve this goal in a large variety of mouse models. We analyzed the literature and performed experiments in three independent institutions, showing that neutrophil depletion using the anti-Ly6G ab (clone 1A8) is not a reliable method in C57BL/6J mice as even at high doses the anti-Ly6G might only be used in mice younger than 12 weeks of age where it leads to a partial depletion (70%). The reproducibility of our observations in mice from different origins strongly suggest that resistance to Ly6G ab-mediated neutrophil depletion is linked with mouse age even if the impact of the microbiota and chronic inflammation could also be important^27^. Understanding why anti-Ly6G treatment works in FVB/N and BALB/c but not in C57BL/6J mice remains an important question. Our observations might highlight differences in hematopoiesis and/or efficiency of the antibody dependent phagocytosis (ADCP) or cell cytotoxicity (ADCC) between mouse strains. Similarly, the fact that old C57BL/6J mice are more resistant to anti-Ly6G treatment might be linked with age-related NK cell functional decay^28^ that could be associated with reduced ADCP, but this phenomenon is usually observed in mice much older than 12 weeks of age. By using Catchup^IVM-red^ mice we further demonstrate that it is not possible to validate neutrophil depletion in anti-Ly6G treated mice using the same ab for flow cytometry. Other abs are commercially available, such as anti-Ly6C, anti-Ly6B or anti-S100A9 abs. As an alternative to ab treatment, we have also generated Catchup^DTR^ mice but neutrophils were unfortunately not sensitive to DT. Two main hypotheses might explain neutrophil resistance to DT in this Ly6G driven DTR expression model: 1) DTR is not expressed enough at the neutrophil membrane, 2) knowing that most of the cytotoxic activity of DT being largely dependent on protein synthesis inhibition^29^, it is possible that early differentiated Ly6G^+^ neutrophils do not require anymore protein synthesis to reach the periphery and to survive for 12 hours^30^. Nevertheless, the Catchup^DTR^ model could be relevant as protein synthesis might be important for neutrophil function in tissues. Interestingly, these results are contrasting with data from MRP8^DTR^ mice where neutrophils, together with a monocyte sub-population, disappear from the circulation upon DT treatment. This might suggest that MRP8 is expressed earlier than Ly6G during neutrophil biogenesis, at a stage of differentiation where progenitors remain sensitive to DT.

Although anti-Ly6G fails to deplete neutrophils in C57BL/6 mice, we are not claiming that it has no impact on neutrophil function as it has helped to identify numerous processes where neutrophils play a central role. We report here that treating mice with neutrophil depleting agents leads to the mobilization of stem cells, progenitors and neutrophil pools in the BM. This observation is supported by the fact that neutrophil phagocytosis in the BM is a central mechanism regulating HSC niches through inhibition of CXCL12 production by reticular cells^11^. Furthermore, neutrophil homeostasis is sensed in the gut by CD169^+^ macrophages in a process governing the IL23/IL17/G-CSF axis, which is pivotal in the regulation of granulopoiesis in the BM^7,10^. Altogether, these results suggest that in C57BL/6J mice neutrophil depletion is impaired by exacerbated neutropoiesis. On the other hand, treating these mice with anti-Gr1 ab leads to profound neutrophil depletion suggesting that the celerity of neutrophil clearance is the limiting parameter in anti-Ly6G treated mice. As the anti-Ly6G ab has been proposed to deplete neutrophils though a mechanism dependent on macrophages^31^, we were able to establish neutrophil clearance by combining anti-Ly6G with a secondary ab^32^. This approach is to date the most efficient and specific method to deplete these cells even when compared to genetically engineered mice allowing LysM, MRP8 or Ly6G-based conditional DTR expression, strategies which show lack of specificity (only 80% of the target cells are neutrophils)^14,17^ or absence of effect on the number of circulating cells (this study).

The most obvious impact of neutrophil depletion on the immune signature was an increase in both AM and IM proportions in the tumor mass associated with a strong reduction of Treg infiltration and amplification. IM showed a lower expression of tumor-promoting genes such as *Mmp9*^33^ and immunomodulatory mediators (*Arg1, Nos2*). Hence, neutrophil infiltration might link some of the alterations reported on lung tumor patients’ AMs^34–36^. Finally, treating KP mice with anti-Ly6G ab alone failed to demonstrate any effect on tumor growth and the immune signature. In our previous study^20^ we demonstrated that depleting neutrophils with anti-Gr1 ab sensitizes tumors to anti-PD1 immunotherapy. Furthermore, a retrospective study suggested that lung cancer patients’ survival was lower following immunotherapy if they had received an antibiotic treatment before^37^. Our work raises the hypothesis that neutrophil interference with macrophage function ultimately contributes to immunosuppressive T cell response with increased Treg amplification in lung tumors. This is an interesting perspective that will require in-depth analyses. A mechanism through which neutrophils could dampen macrophages’ pro-inflammatory functions and potential anti-tumor activity might be directly linked to aged neutrophil phagocytosis by these cells, a phenomenon leading to suppression of pro-inflammatory responses by macrophages in multiple tissues^39^.

To conclude, our work illustrates critical limitations inherent to the use of anti-Ly6G ab for neutrophil depletion. While we urge the scientific community to systematically use and report appropriate methods to validate neutrophil depletion, for the last ten years, it is possible that some effects reported from anti-Ly6G treatment in C57BL/6 mice in various models are linked to aged neutrophils rather than total neutrophil biological activity. Our work proposes an alternative method for specific and efficient neutrophil depletion.

## Online Methods

### Mice importation and housing conditions

All mouse experiments from the Ecole Polytechnique Fédérale de Lausanne were performed with the permission of the Veterinary Authority of the Canton de Vaud, Switzerland (license number VD2391). Mouse experiments at Massachusetts General Hospital were performed according to approved IACUC guidelines. Wild type C57BL/6 mice were purchased from Jackson Laboratory. KP1.9 tumors were injected intravenously as previously described^40^. Mouse experiments performed at the Cancer Research Center of Lyon at the Center Léon Bérard, Lyon, France were approved by the local Animal Ethic Evaluation Committee (CECCAPP: C2EA-15) and authorized by the French Ministry of Education and Research. Animals were maintained in a specific pathogen free (SPF) animal facility AniCan platform.

### Management of the KP mouse model of lung tumor

The tumors were initiated upon *in vivo* transduction of lung epithelial cells with a viral vector delivering Cre recombinase to activate oncogenic *Kras^G12D^* and delete *p53*. Twelve-to-fourteen-week-old mice were instilled intratracheally with 1500 Cre-active lentiviral units. Neutrophil depletion was performed on week 25 after tumor initiation. To measure tumor volume, mice were anaesthetized using isoflurane and maintained under anesthesia during the scanning procedure. Lungs were imaged with a μCT (Quantum FX, PerkinElmer) at a 50-μm voxel size, with retrospective respiratory gating. Individual tumor volumes were measured and calculated using the Analyze software (PerkinElmer).

### Flow cytometry and cell sorting

For 17 colors flow cytometry see supplementary method part b.

All acquisitions were performed using the LSRII SORP (Becton Dickinson), a 5-laser and 18-detector analyzer at the EPFL Flow Cytometry Core Facility. Data analyses were performed using FlowJo X (FlowJo LLC ©). True-count flow cytometry experiment were performed on 30μl of total blood directly stained with antibodies wash into 2ml of PBS then complemented with 25μl of countBright absolute counting beads (C3690 Invitrogen) and resuspended into 2 ml of PBS before acquisition on a Attune NxT cytometer (ThermoFisher).

Sorting of alveolar (DAPI-CD45^+^CD11b^−^F4/80^+^SiglecF^+^) and interstitial macrophages (DAPI^−^CD45^+^CD11b^+^F4/80^+^SiglecF^−^) were performed using FASC-ARIA-II and FACS-ARIA-Fusion on pre-enriched CD45^+^ cells from tumor single cell suspensions using magnetic cell sorting (CD45 MicroBeads, 130-052-301 and AutoMACS device from Miltenyi). Neutrophils were isolated using Miltenyi Ly6G purification microbeads (103-092-332).

Fluorescence activated cell sorting was performed using the MoFlow ASTRIOS EQ cell sorter. Before sorting, immune cells were enriched using CD45 magnetic isolation. Neutrophils (CD11b^+^ Ly6G^+^), monocytes (CD11b^+^, CD11c^−^, F4/80^−^, CD3^−^, B220^−^), T cells (CD3^+^), B cells (B220^+^, CD11c^−^), macrophages (CD11b^+/int^, F4/80^+^) and DCs (CD11c^+^ F4/80^−^, CD11b^+/int^, Ly6G^−^) were sorted simultaneously among CD45^+^ DAPI^−^ viable cells from the same sample.

## Supporting information

## Acknowledgements

We thank the EPFL SV Flow Cytometry Core Facilities for access to instruments and technical help for cell sorting. This work was supported by the Swiss National Science Foundation (310030_179324), the ISREC Foundation, the Chercher et Trouver Foundation, the Nuovo Soldati Foundation (to GB), the MGH ECOR (Executive Committee on Research) Tosteson Postdoctoral Fellowship (to CP), a postdoctoral fellowship from the Fondation pour la Recherche Médicale (SPF20160936218) (to JM), the French National Cancer Institute INCa 2014-093, INCa 2017-158, the Ligue contre le cancer, and the LABEX DevWeCan.

## Author contributions

JF conceptualized the study, performed experiments and wrote the manuscript. GB and MCV provided results of blood smears analysis from Ly6G and Gr1 ab treated mice. AG performed mouse necropsy, flow cytometry staining, acquired most of the data and wrote the supplementary method section. PBA participated in mouse treatment and KP mouse management. JM, NBV, CC, CE, CP and MJP produced data on neutrophil depletion using anti-Ly6G ab done in their respective laboratory. MG generated and provided Catchup mice. JV performed immunostaining against HB-EGF on purified neutrophils and contributed to KP mouse management. EM supervised the study and edited the manuscript. All authors analyzed the data.

## References

1. Phillipson, M. & Kubes, P. The neutrophil in vascular inflammation. Nat. Med. 17, 1381–1390 (2011).

2. Shen, M. et al. Tumor-Associated Neutrophils as a New Prognostic Factor in Cancer: A Systematic Review and Meta-Analysis. PLoS ONE 9, (2014).

3. Lee, K. H. et al. Neutrophil extracellular traps (NETs) in autoimmune diseases: A comprehensive review. Autoimmun. Rev. 16, 1160–1173 (2017).

4. Kaplan, M. J. Role of neutrophils in systemic autoimmune diseases. Arthritis Res. Ther. 15, 219 (2013).

5. Reber, L. L. et al. Neutrophil myeloperoxidase diminishes the toxic effects and mortality induced by lipopolysaccharide. J. Exp. Med. jem.20161238 (2017). doi:10.1084/jem.20161238

6. Wilgus, T. A., Roy, S. & McDaniel, J. C. Neutrophils and Wound Repair: Positive Actions and Negative Reactions. Adv. Wound Care 2, 379–388 (2013).

7. Casanova-Acebes, M. et al. Neutrophils instruct homeostatic and pathological states in naive tissues. J. Exp. Med. jem.20181468 (2018). doi:10.1084/jem.20181468

8. Saverymuttu, S. H., Peters, A. M., Keshavarzian, A., Reavy, H. J. & Lavender, J. P. The kinetics of 111Indium distribution following injection of 111Indium labelled autologous granulocytes in man. Br. J. Haematol. 61, 675–685 (1985).

9. Suratt, B. T. et al. Neutrophil maturation and activation determine anatomic site of clearance from circulation. Am. J. Physiol.-Lung Cell. Mol. Physiol. 281, L913–L921 (2001).

10. Stark, M. A. et al. Phagocytosis of Apoptotic Neutrophils Regulates Granulopoiesis via IL-23 and IL-17. Immunity 22, 285–294 (2005).

11. Casanova-Acebes, M. et al. Rhythmic Modulation of the Hematopoietic Niche through Neutrophil Clearance. Cell 153, 1025–1035 (2013).

12. Lee, P. Y., Wang, J.-X., Parisini, E., Dascher, C. C. & Nigrovic, P. A. Ly6 family proteins in neutrophil biology. J. Leukoc. Biol. 94, 585–594 (2013).

13. Daley, J. M., Thomay, A. A., Connolly, M. D., Reichner, J. S. & Albina, J. E. Use of Ly6G-specific monoclonal antibody to deplete neutrophils in mice. J. Leukoc. Biol. 83, 64–70 (2008).

14. Clausen, B. E., Burkhardt, C., Reith, W., Renkawitz, R. & Förster, I. Conditional gene targeting in macrophages and granulocytes using LysMcre mice. Transgenic Res. 8, 265–277 (1999).

15. Faust, N., Varas, F., Kelly, L. M., Heck, S. & Graf, T. Insertion of enhanced green fluorescent protein into the lysozyme gene creates mice with green fluorescent granulocytes and macrophages. Blood 96, 719–726 (2000).

16. Lagasse, E. bcl-2 inhibits apoptosis of neutrophils but not their engulfment by macrophages. J. Exp. Med. 179, 1047–1052 (1994).

17. Abram, C. L., Roberge, G. L., Hu, Y. & Lowell, C. A. Comparative analysis of the efficiency and specificity of myeloid-Cre deleting strains using ROSA-EYFP reporter mice. J. Immunol. Methods 408, 89–100 (2014).

18. Hasenberg, A. et al. Catchup: a mouse model for imaging-based tracking and modulation of neutrophil granulocytes. Nat. Methods 12, 445–452 (2015).

19. Madisen, L. et al. A robust and high-throughput Cre reporting and characterization system for the whole mouse brain. Nat. Neurosci. 13, 133–140 (2010).

20. Faget, J. et al. Neutrophils and Snail Orchestrate the Establishment of a Pro-tumor Microenvironment in Lung Cancer. Cell Rep. 21, 3190–3204 (2017).

21. Engblom, C. et al. Osteoblasts remotely supply lung tumors with cancer-promoting SiglecFhigh neutrophils. Science 358, eaal5081 (2017).

22. Brüggemann, M. Evolution of the rat immunoglobulin gamma heavy-chain gene family. Gene 74, 473–482 (1988).

23. Buch, T. et al. A Cre-inducible diphtheria toxin receptor mediates cell lineage ablation after toxin administration. Nat. Methods 2, 419–426 (2005).

24. Tan, S. Y. S. & Krasnow, M. A. Developmental origin of lung macrophage diversity. Dev. Camb. Engl. 143, 1318–1327 (2016).

25. Roberts, L. M. et al. Depletion of alveolar macrophages in CD11c diphtheria toxin receptor mice produces an inflammatory response. Immun. Inflamm. Dis. 3, 71–81 (2015).

26. Soroosh, P. et al. Lung-resident tissue macrophages generate Foxp3^+^ regulatory T cells and promote airway tolerance. J. Exp. Med. 210, 775–788 (2013).

27. Deshmukh, H. S. et al. The microbiota regulates neutrophil homeostasis and host resistance to Escherichia coli K1 sepsis in neonatal mice. Nat. Med. 20, 524–530 (2014).

28. Fang, M., Roscoe, F. & Sigal, L. J. Age-dependent susceptibility to a viral disease due to decreased natural killer cell numbers and trafficking. J. Exp. Med. 207, 2369–2381 (2010).

29. Kageyama, T. et al. Diphtheria Toxin Mutant CRM197 Possesses Weak EF2-ADP-ribosyl Activity that Potentiates its Anti-tumorigenic Activity. J. Biochem. (Tokyo) 142, 95–104 (2007).

30. Evrard, M. et al. Developmental Analysis of Bone Marrow Neutrophils Reveals Populations Specialized in Expansion, Trafficking, and Effector Functions. Immunity 48, 364–379.e8 (2018).

31. Bruhn, K. W., Dekitani, K., Nielsen, T. B., Pantapalangkoor, P. & Spellberg, B. Ly6G-mediated depletion of neutrophils is dependent on macrophages. Results Immunol. 6, 5–7 (2015).

32. Goldschmidt, T. J., Holmdahl, R. & Klareskog, L. Depletion of murine T cells by in vivo monoclonal antibody treatment is enhanced by adding an autologous anti-rat κ chain antibody. J. Immunol. Methods 111, 219–226 (1988).

33. Mehner, C. et al. Tumor cell-produced matrix metalloproteinase 9 (MMP-9) drives malignant progression and metastasis of basal-like triple negative breast cancer. Oncotarget 5, 2736–2749 (2014).

34. Pouniotis, D. S., Plebanski, M., Apostolopoulos, V. & McDonald, C. F. Alveolar macrophage function is altered in patients with lung cancer. Clin. Exp. Immunol. 143, 363–372 (2006).

35. Dewhurst, J. A. et al. Characterisation of lung macrophage subpopulations in COPD patients and controls. Sci. Rep. 7, (2017).

36. Bazzan, E. et al. Dual polarization of human alveolar macrophages progressively increases with smoking and COPD severity. Respir. Res. 18, (2017).

37. Derosa, L. et al. Negative association of antibiotics on clinical activity of immune checkpoint inhibitors in patients with advanced renal cell and non-small-cell lung cancer. Ann. Oncol. 29, 1437–1444 (2018).

38. Le Noci, V. et al. Modulation of Pulmonary Microbiota by Antibiotic or Probiotic Aerosol Therapy: A Strategy to Promote Immunosurveillance against Lung Metastases. Cell Rep. 24, 3528–3538 (2018).

39. A-Gonzalez, N. et al. Phagocytosis imprints heterogeneity in tissue-resident macrophages. J. Exp. Med. 214, 1281–1296 (2017).

40. Pfirschke, C. et al. Immunogenic Chemotherapy Sensitizes Tumors to Checkpoint Blockade Therapy. Immunity 44, 343–354 (2016).

