## Supplementary material for "Efficient and specific Ly6G^+^ cell depletion: A change in the current practices toward more relevant functional analyses of neutrophils"

Supplementary figure legends

**Figure S1: Inefficiency of the anti-Ly6G depletion is observed across institutions**

a) Histogram shows percentage of neutrophils in spleen, lung and BM from lung tumor transplanted

C57BL/6J mice treated with 500μg of anti-Ly6G (n=7) or control ab (Ctr n=6) every two/three days over thirteen days. Tail vein KP derived lung cancer cells transplantation was done twenty days before anti-Ly6G treatment initiation. Raw flow cytometry files were provided by M. Pittet's lab (Massachusetts General Hospital Research Institute/Harvard Medical School, USA).

b) Histogram shows percentage of neutrophils in blood before and at day one or day six of treatment with 150μg of anti-Ly6G ab injected daily in C57BL/6J mice produced in house (black dots) or imported from Charles River (red dots). Raw flow cytometry files were provided by C. Caux's lab (INSERM, CLB/Centre Leon Bérard, France). This experiment is representative of 13 independent ones, where anti-Ly6G was tested at different concentrations (from 100μg to 800μg) and at different frequency (daily to weekly).

a and b) neutrophils were identified among $CD45^+$ cells as $CD11b^+Ly6C^{int}$ cells. ** $p<0.01$ from Mann-Whitney test, error bars represent SEM.

c) Curves show the mean fluorescence intensity (MFI) of the indicated abs on $Tomato^+CD11b^+$ neutrophils over an antibody titration assay ranging from 25ng/ml to 20μg/ml and performed on blood samples from $Catchup^{IVM\text{-}red}$ mice treated with 100μg of control (Ctr: circle) anti-Ly6G (square) or anti-Gr1 (triangle) abs, 24 hours before blood sampling.

**Figure S2: anti-Ly6G treatment as well as efficient neutrophil depletion leads to the disappearance of the mature neutrophil pool present in the BM**

a) Gating strategy used to identify GMP cells in BM.

b) Flow cytometry plot showing the proportion of mature neutrophils ($CXCR2^+CD62L^+$) in BM from control (n=4), anti-Ly6G alone (n=5) or anti-Ly6G plus anti-Rat-IgGk (n=5) treated mice. Histograms gives the percentage of mature (left) and immature (right) neutrophils among $CD11b^+S100A9^+$ cells in the BM. * $p<0.05$ from Mann-Whitney test, error bars represent SEM.

**Figure S3: Ly6G$^{+}$ cells are resistant to DT and neutralization of anti-Ly6G ab due accumulation of anti-rat-IgGk ab in vivo can abrogate neutrophil depletion**

a) HB-EGF staining (brown) on sorted neutrophils from Catchup$^{DTR}$ (Ly6G$^{KI/WT}$; ROSA26$^{LSL\text{-}HB\text{-}EGF}$) or littermate BMs (Ly6G$^{WT/WT}$; ROSA26$^{LSL\text{-}HB\text{-}EGF}$).

b) Flow cytometry plots showing the staining and gating strategy to monitor neutrophil apoptosis in vitro after 12 hours of culture in presence of DT from 0 to 1μg/ml. Histogram represents the results of the same experiment done in triplicate. Error bar show SD.

c) Dot plot shows the depletion efficiency (%) at the indicated time points in regard to the proportion of neutrophils at day 0 during combination treatment (anti-Ly6G and anti-rat-IgGk as in figure 3). Identical symbol represents the same mouse along the time course.

d) Flow cytometry plot showing the gating strategy used to identify neutrophils and histograms give the fluorescence intensity of the anti-Ly6G ab on neutrophils in anti-Ly6G alone or combination treatment as in c). The numbers of remaining neutrophils per ml of blood from true-count flow cytometry are indicated. Colors of the histograms indicate the time points of blood sampling.

e) Scatter plots evaluating the correlation between the number of remaining neutrophils and the MFI of anti-Ly6G staining in the same experiment. * $p<0.05$ from spearman correlation test.

**Figure S4: neutrophil depletion associates with partial modification of tumor macrophage behavior**

a) Percentage of neutrophils among blood circulating cells in KP mice bearing well established tumors and treated with ctr + anti-rat, anti-Ly6G or anti-Ly6G plus anti-rat over 11 days (as in figure 4b), n=5.

b) KP mice were treated with 200μg of anti-Gr1 ab (n=10) or control ab (n=11) every two days over three weeks and macrophage percentage was determined among immune cells as total $F4/80^{+}$ cells.

c) Histograms represents the relative mRNA expression of the indicated genes on sorted AM ($CD45^{+}CD11b^{-}SiglecF^{+}F4/80^{+}$) and IM ($CD45^{+}CD11b^{+}F4/80^{+}$) from KP tumors following control (ctr-rat) and combination treatment (Ly6G-rat).

d) Histogram shows the percentage of $Ki67^{+}$ cells among Treg ($CD45^{+}CD4^{+}CD3^{+}Foxp3^{+}$) in tumor from mice that received ctr + anti-rat (n=8) anti-Ly6G (n=7) or anti-Ly6G + anti-rat (n=10) treatment over 12 days.

e) Percentage of Treg among $CD4^{+}$ T cells identified as in a) in tumors from mice treated over three weeks with anti-Gr1 or control ab.

Error bars indicate SEM, * $p<0.05$, ** $p<0.01$, *** $p<0.001$ from Mann-Whitney test

Efficient and specific Ly6G+ cell depletion: A change in the current practices toward more relevant functional analyses of neutrophils

Supplementary methods

**a) Neutrophil depletion**

This protocol describes the method by which efficient and specific neutrophil depletion can be achieved in C57BL/6 mice. Two widely used strategies in the literature identified for this purpose were tested for their efficacy and specificity: the use of anti-Gr1 (clone: RB6-5C8), or anti-Ly6G (clone: 1A8) antibody. Although administration of the anti-Gr1 Ab leads to an extended drop in the proportion of blood neutrophils, its binding to both Ly6G and Ly6C antigens leads to unspecific depletion of neutrophils, and Ly6C$^{+}$CD8$^{+}$ T cells and monocytes. On the other hand, the anti-Ly6G Ab, which exclusively binds to neutrophils, fails to reduce their number in the circulation of C57BL/6 mice. We validated a combination of the neutrophil-specific antibody, anti-Ly6G, together with the secondary anti-Rat IgG2aκ light chain (clone: MAR18.5) ab that leads to an efficient neutrophil depletion.

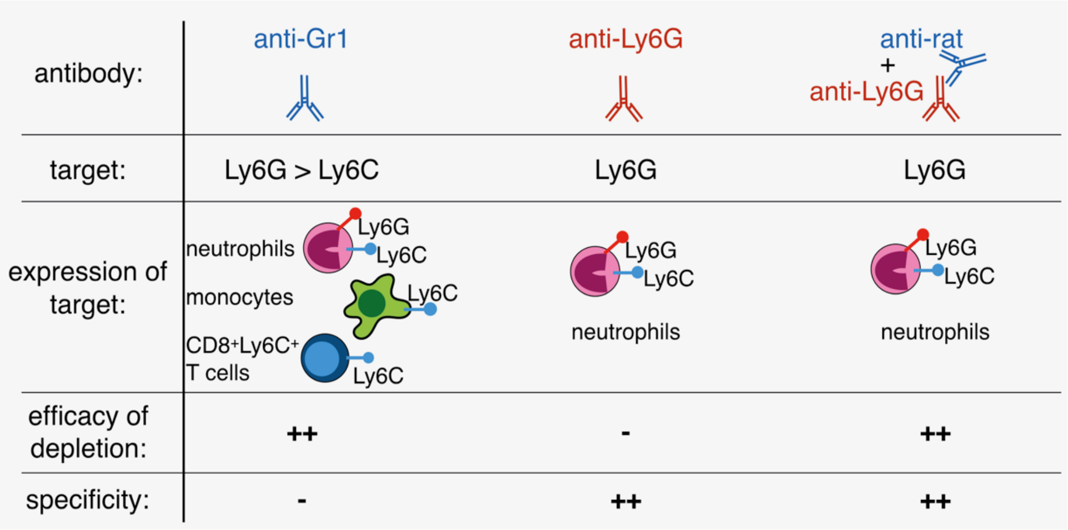


**Methods**

For short-term experiments (max. five days, from Day 1 to 5), highly efficient neutrophil depletion is achieved in C57BL/6J mice by intraperitoneal (IP) injection with 100µg of anti-Ly6G antibody (clone: 1A8) at days 0, 2, 5 and 100µg of anti-Rat IgGk (clone: MAR18.5) at days 1and 3. To evaluate neutrophil depletion, blood sampling (30-50µl) from lateral tail vein must be performed before any ab injection, at day 0, day 2 and day 5.

For long-term depletion (15 days and longer), this strategy is limited by the accumulation of anti-rat-IgGκ ab in the mouse serum, leading to neutralization of the anti-Ly6G ab injected to target neutrophils. Hence, careful monitoring of the number of circulating neutrophils (identified as CD11b$^{+}$Ly6C$^{int}$ cells) must be performed every two days from day 5 using anti-CD11b, anti-Ly6C plus anti-Ly6G abs for flow cytometry. The Mean Fluorescence Intensity (MFI) of the Ly6G staining on CD11b$^{+}$Ly6C$^{int}$ must be assessed to validate the binding on residual neutrophils of the anti-Ly6G ab injected *in vivo*. If anti-Ly6G antigen is accessible for flow cytometry staining with fluorophore-coupled anti-Ly6G ab, it means that the anti-Ly6G ab injected *in vivo* had been neutralized by an excess of anti-rat-IgGκ ab. Conversely, if residual neutrophils appear negative for anti-Ly6G staining in flow cytometry analyses, this suggests that anti-rat-IgGκ ab concentration is too low in the animals.

We have validated the experimental procedure described below.

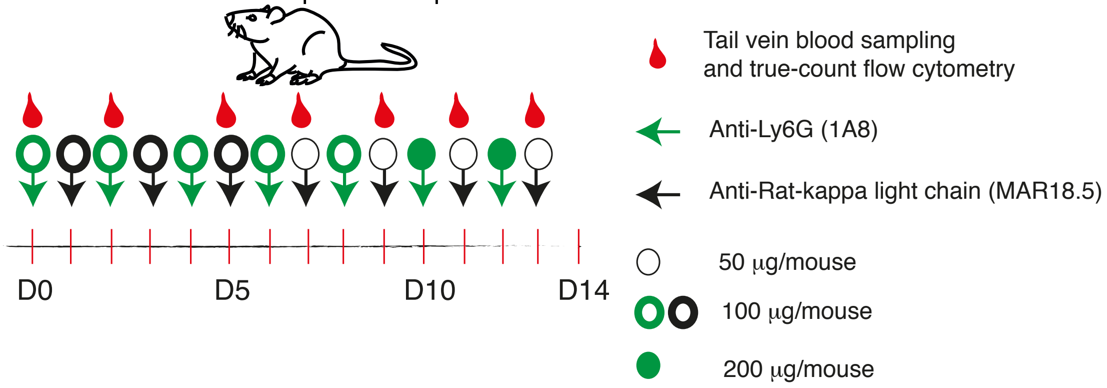


For two weeks depletion, mice are treated with 100μg of anti Ly6G ab IP at day 0, 2, 4, 6, 8 then 200μg at day 10 and 12. Mice receive IP injection of 100μg of anti-rat-IgGκ at day 1, 3, 5 then 50μg at day 7, 9, 11 and 13.

For all these combination treatments we concluded that the most relevant control is to use the irrelevant ab clone 2A3 in place of the anti-Ly6G ab. It is important to keep treating the mice with the anti-rat-IgGκ ab following the exact same procedure.

The main limitation of this approach is linked to the usage of the anti-rat-IgGκ as it requires constant monitoring of neutrophil proportion and anti-Ly6G binding. Furthermore, this strategy is not compatible with any treatment combination that include other rat-IgGκ antibodies such as anti-PD1 ab as it will also lead to depletion of PD1 positive cells for example.

<u>Antibodies:</u>

| **Target** | **Clone** | **Source** |
|---|---|---|
| Control ab | 2A3 | BioXcell |
| Anti-Ly6G | 1A8 | BioXcell |
| Anti-rat IgGκ | MAR18.5 | BioXcell |

**b) Extracting the immune signature of KP tumors**

This method relies on careful dissection of individual tumors. To compare different conditions it is important to measure tumor weight and compare similar size lesions as the immune signature is strongly impacted by the ratio between volume and surface of the tumors.

1) Mice were sacrificed using pentobarbital. Avoid using asphyxia on animals displaying advance disease stages as it can cause lung hemorrhage decreasing the quality of the results. Intra-cardiac terminal blood sampling is recommended to reduce blood contamination.

2) Lung tumors are resected using a stereoscope; it is critical to avoid contamination of the samples with surrounding lung tissues. This method is working on small (5mg) to very big (300mg) tumors. Individual tumors are weighed then mechanical and enzymatic dissociations are performed.

2.1) Tumors are placed in 100ml of DMEM for 20mg of tissue then chopped with chisels in small pieces. Supernatants can be harvested then centrifuged, filtered and frozen for further analysis of soluble factors.

2.2) Tissue is then suspended in 2.5 ml of digestion mix containing 9.4mg/ml collagenase I and 0.25g/ml DNase I in DMEM. The enzymatic digestion is performed using GentelMACS Octo-dissociator or alternatively in 24 well plate at 37°C for 45 min under magnetic agitation.

2.3) Single cell suspension are filtered in through a 70µm cell strainer in 7.5 ml of DMEM containing 5% fetal calf serum.

3) Flow cytometry staining is performed on a maximum of 20mg of tissues in 96 V bottom plate.

3.1) Single cell suspensions produced in 2.3 are centrifuged (450rcf/7min) then cells are suspended considering the weight of the lesions measured in point 2.1 in PBS: 100µl/20mg (for tumor inferior to 20mg use a minimal volume of 100µl). 100µl of each sample is then transferred into the 96 well plate then the pate is centrifuged 450rcf/4min. Of note, all samples will be processed simultaneously in the 96 well plate hence it is important to consider including extra-wells for compensation controls and fluorescence controls / isotype fluorescence minus one (FMO) controls that will be used to set the cytometer and analyses flow cytometry files.

3.2) Supernatant is removed then cell pellets are resuspended in 50µl of PBS containing the fixable cell viability dye (Live and Dead Blue life technologies) and Fc-Block reagent dilution 1/100. Then the pellet is stored at 4°C for 20 min.

3.3) Prepare 50µl of ab mix per well for membrane staining at a 2X concentration (ex: ab used at a final dilution 1/200 should be prepared at a 1/100 dilution in this mix) in PBS containing 2% bovine serum albumin (BSA) or 2% FCS (FACS buffer). Then add 50µl of mix per well (volume per well is now 100µl) and incubate 15min at 4°C.

3.4) Add 100µl of FACS buffer, centrifuge the plate 450rcf/4min then repeat washing with 200ml of FACS buffer. Remove FACS buffer then fix the cells with 50µl of 1X Perm and Fix solution of the FoxP3 / transcription factor buffer set from eBioscience (00-5523-00) for 25min at 4°C. Wash by adding 150µl of FACS buffer and centrifugation 450rcf/4min then replace supernatant with 200µl of FACS buffer. Once covered with plastic film the plate can be kept at 4°C for one week protected from light.

3.5) Perform intra-nuclear / cytoplasmic staining the day of flow cytometry acquisition. Centrifuge the plate 450rcf/4min, remove FACS buffer then resuspend the cells in 200µl of Perm solution 1X of the FoxP3 / transcription factor buffer set from eBioscience. Centrifuge the plate again 450rcf/4min and remove supernatant then proceed to intra-cellular staining in a volume of 100µl of ab mix and incubate 20min 4°C. Add 100ml of Perm buffer 1x centrifuge 450rcf/4min then repeat washing with 200µl of perm buffer then with 200µl of FACS buffer. Resuspend the cells in 150µl of FACS buffer, transfer each well in a micronic tube for flow cytometer acquisition. Of note, to increase the stability during data acquisition, an extra filtration step at the end of the staining is recommended.

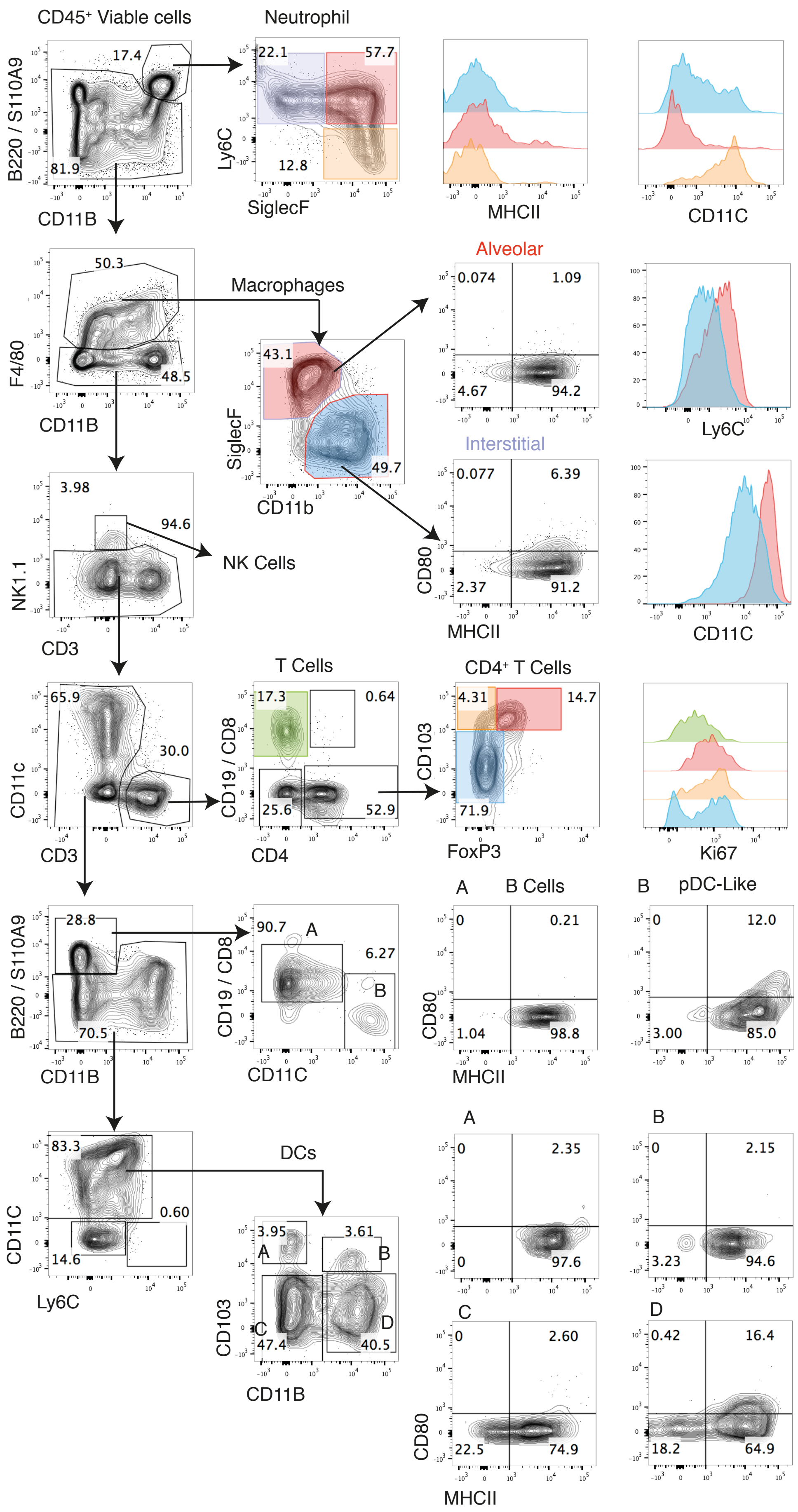
CD45+ Viable cells
Neutrophil
B220 / S110A9
CD11B
Ly6C
SiglecF
MHCII
CD11C
Alveolar
Interstitial
Macrophages
F4/80
SiglecF
CD11b
CD80
Ly6C
NK Cells
NK1.1
CD3
T Cells
CD4+ T Cells
CD11c
CD19 / CD8
CD4
CD103
FoxP3
Ki67
B Cells
pDC-Like
CD11C
DCs
CD11B
A
B
C
D

17 colors flow cytometry panel and analysis:

| Target | Fluorophore | Clone | Source | Identifier |
|---|---|---|---|---|
| anti-FoxP3 | FITC | FJK-16S | Thermofisher | Ref: 53577382 |
| anti-Ly6C | PerCp-Cyanine5.5 | HK1.4 | Biolegend | Ref: 128012 |
| anti-CD11c | Brilliant Violet 450 | N418 | Biolegend | Ref: 117330 |
| anti-CD19/<br>anti-CD8a | Brilliant Violet 510 | 6D5<br>53-6.7 | Biolegend<br>BD Horizon | Ref: 115546<br>Ref: 563068 |
| anti-Ki67 | Brilliant Violet 605 | 16A8 | Biolegend | Ref: 652413 |
| anti-NK1.1 | Brilliant Violet 660 | PK136 | Biolegend | Ref: 108736 |
| anti-CD11b | Brilliant Violet 710 | M1/70 | Biolegend | Ref: 101241 |
| anti-CD4 | Brilliant Violet 780 | RM4-5 | Biolegend | Ref: 100453 |
| anti-B220/<br>anti-S100A9 | APC | RA3-6B2<br>MRP-14 | Biolegend<br>BD Pharmingen | Ref: 103212<br>Ref: 565833 |
| anti-MHCII | APC700 | M5/114.152 | Biolegend | Ref: 107622 |
| anti-CD80 | APC-Cyanine7 | 16-10A1 | Biolegend | Ref: 104740 |
| anti-CD103 | PE | 2E7 | eBioscience | Ref: 12-1031-82 |
| anti-SiglecF | PE 610 | REA798 | Miltenyi | Ref: 130-112-172 |
| anti-CD3 | PE-Cyanine5.5` | 145-2C11 | eBioscience | Ref: 35-0031-82 |
| anti-F4/80 | PE-Cyanine7 | BM8 | Biolegend | Ref: 123114 |
| Live and Dead - Blue | NA | NA | Thermofisher | Ref: L34962 |
| anti-CD45 | Brilliant Ultra Violet 660 | 30-F11 | BD Horizon | Ref: 565079 |

**Reagents :**

| Name | Source | Identifier |
|---|---|---|
| Ethylendiaminetetraacetic acid (EDTA) | Axon lab AG | Ref:205-358-3 |
| Fetal Bovine Serum (FBS) | Life Technologies | Ref:10270-106 |
| Bovine Serum Albumin (BSA) | Sigma aldrich | Ref: A7906-100G |
| DMEM | Thermo Fisher | Ref: 41-965-039 |
| Fc-blocking Reagent | Miltenyi | Ref:130-092-575 |

**Tubes and filters:**

| Name | Source | Identifier |
|---|---|---|
| gentleMACS tubes | Miltenyi | Ref: 130-096-334 |
| Corning Falcon Test Tubes with Cell Strainer Snap Cap | Thermo Fisher | Ref:08-771-23 |
| pre-separation filters 70µm | Miltenyi | Ref: 130-095-823 |

**Instruments:**

| Name | Source |
|---|---|
| gentleMACS octo Dissociator | Miltenyi |
| LSRII SORP | Becton Dickinson |
| LSR Fortessa | Becton Dickinson |
