## Supplementary figures and images for "Efficient and specific Ly6G^+^ cell depletion: A change in the current practices toward more relevant functional analyses of neutrophils"

### Supplementary file 2

#### Figure S1

**a**

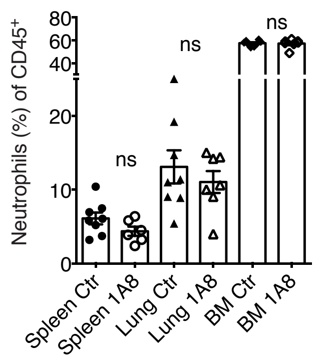


500µg 2-3 times per week
Check Day 13
tumor transplantation 20 days before

**b**

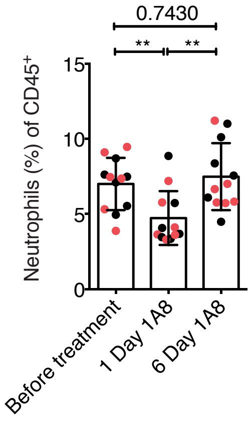


**c**

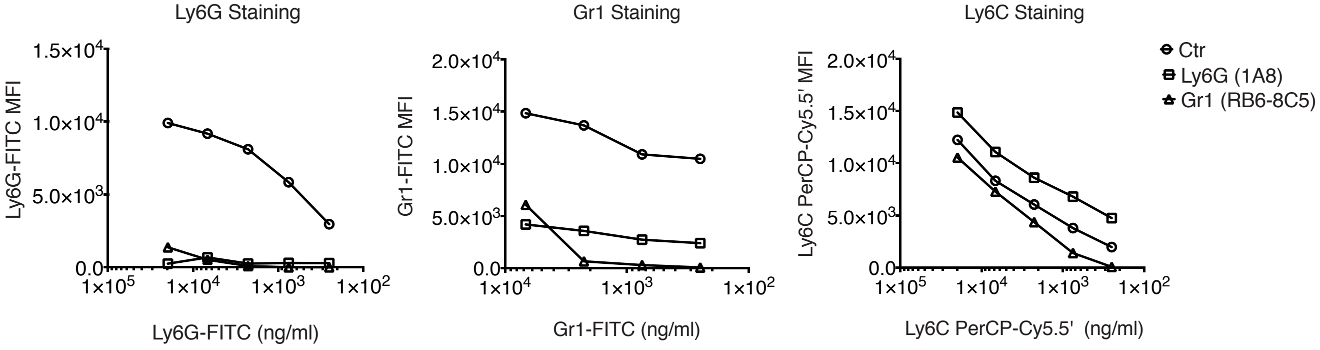

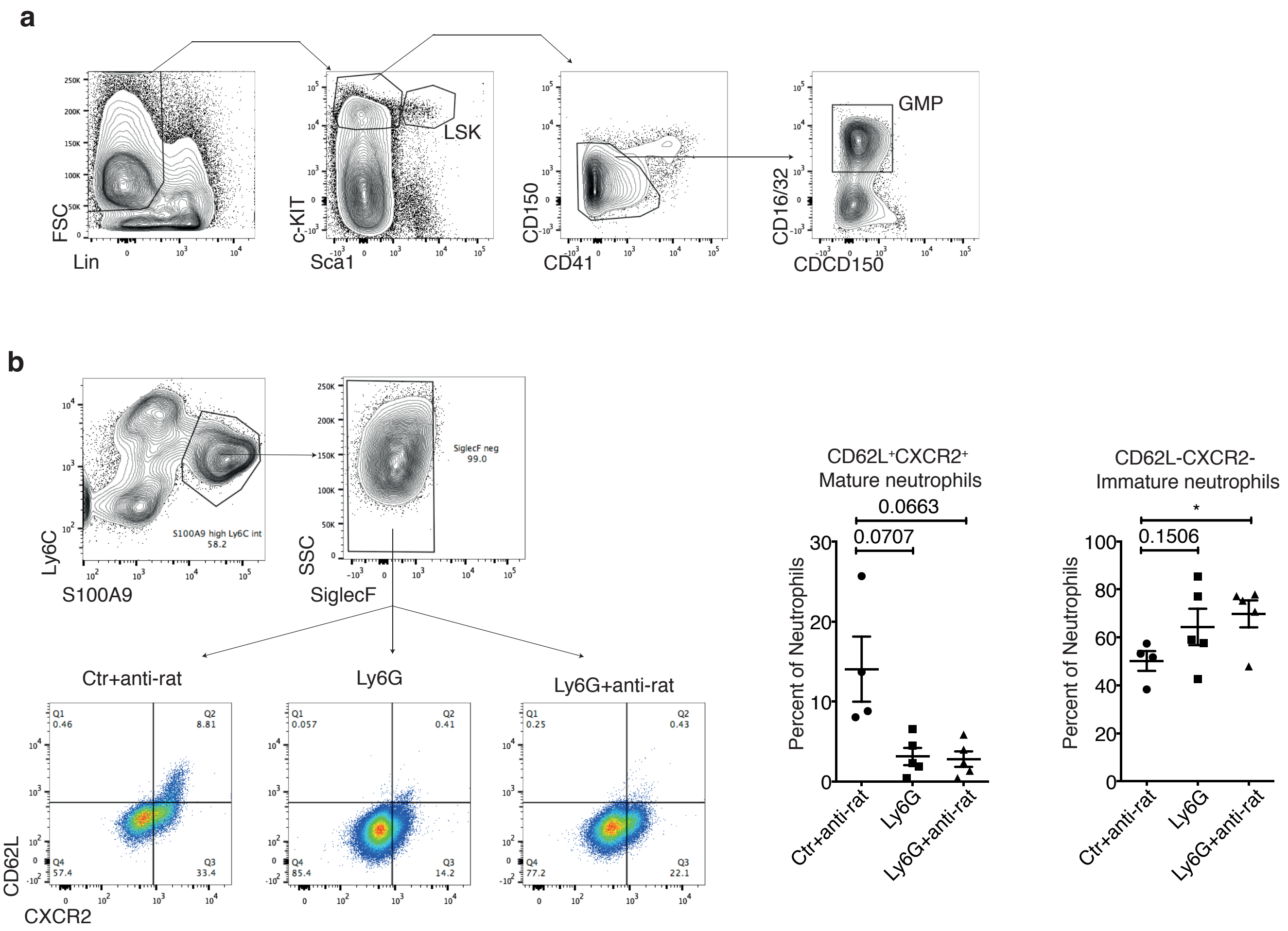

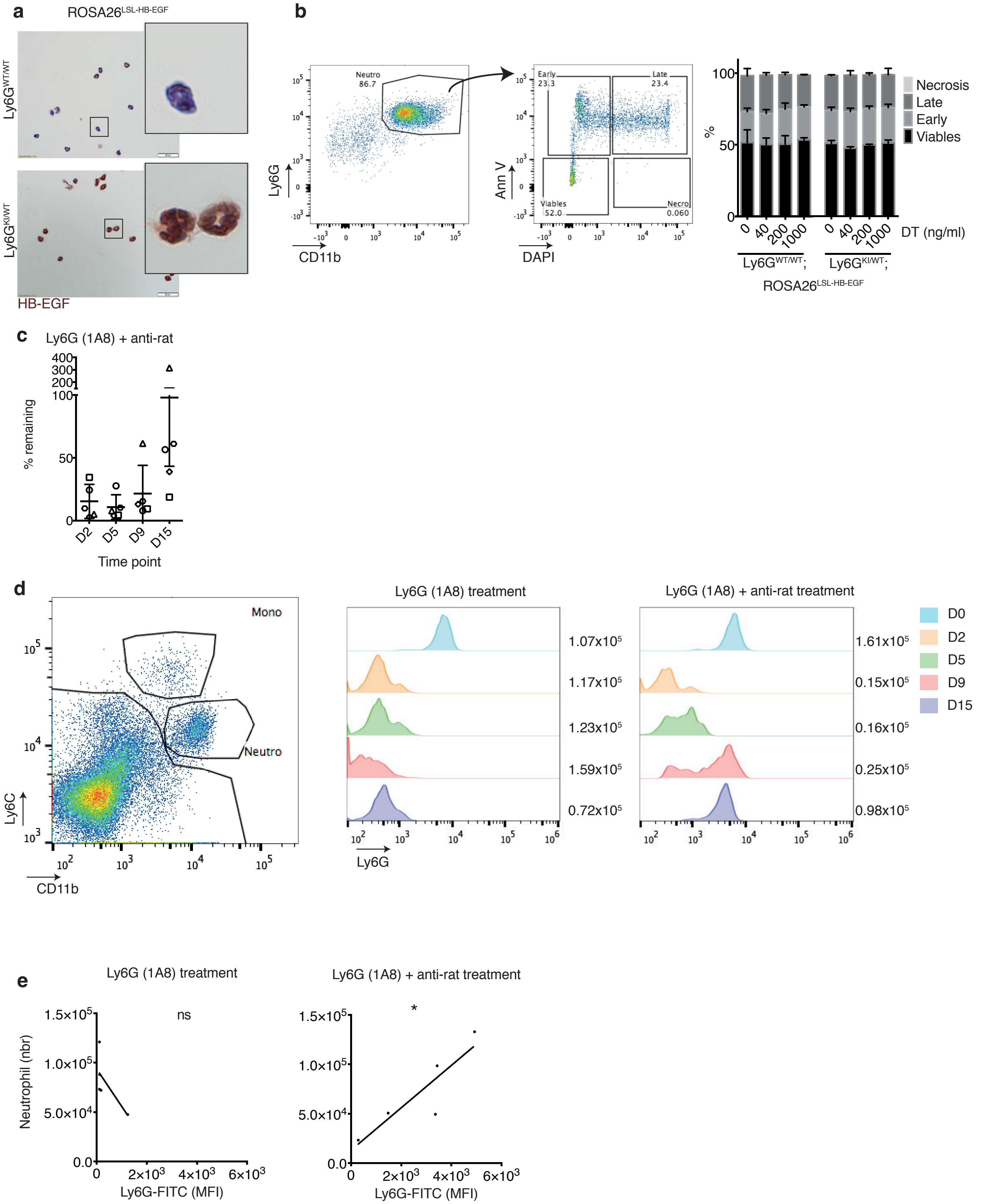

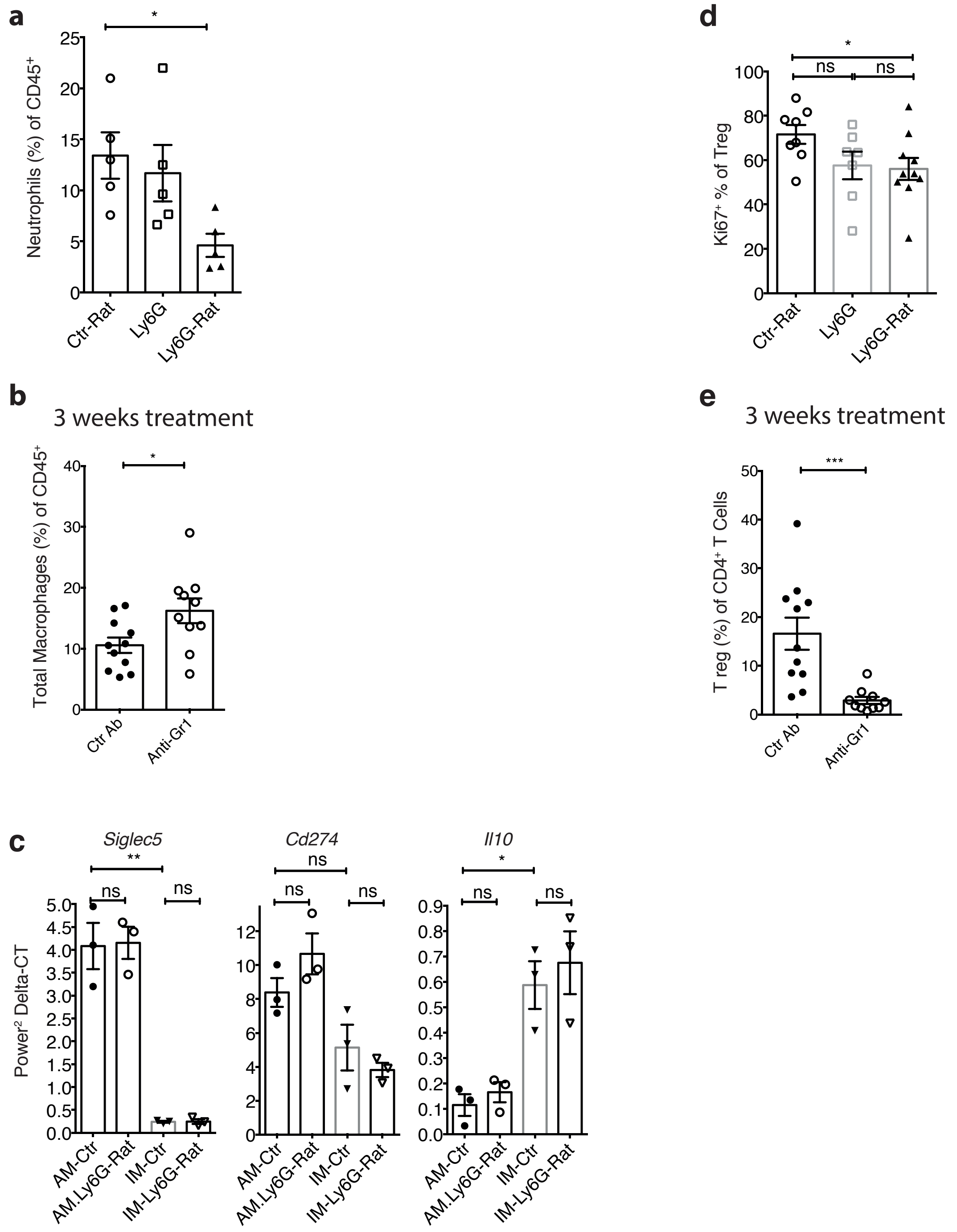
